# Reducing monocarboxylate transporter MCT1 worsens experimental diabetic peripheral neuropathy

**DOI:** 10.1101/2020.02.26.965152

**Authors:** Mithilesh Kumar Jha, Xanthe Heifetz Ament, Fang Yang, Ying Liu, Michael J. Polydefkis, Luc Pellerin, Brett M. Morrison

## Abstract

Diabetic peripheral neuropathy (DPN) is one of the most common complications in diabetic patients. Though the exact mechanism for DPN is unknown, it clearly involves metabolic dysfunction and energy failure in multiple cells within the peripheral nervous system (PNS). Lactate is an alternate source of metabolic energy that is increasingly recognized for its role in supporting neurons. The primary transporter for lactate in the nervous system, monocarboxylate transporter-1 (MCT1), has been shown to be critical for peripheral nerve regeneration and metabolic support to neurons/axons. In this study, MCT1 was reduced in both sciatic nerve and dorsal root ganglia in wild-type mice treated with streptozotocin (STZ), a common model of type-1 diabetes. Heterozygous MCT1 null mice treated with STZ developed a more severe DPN compared to wild-type mice, as measured by greater axonal demyelination, decreased peripheral nerve function, and increased numbness to innocuous low-threshold mechanical stimulation. Given that MCT1 inhibitors are being developed as both immunosuppressive and chemotherapeutic medications, our results suggest that clinical development in patients with diabetes should proceed with caution. Collectively, our findings uncover an important role for MCT1 in DPN and provide a potential lead toward developing novel treatments for this currently untreatable disease.

## Introduction

Diabetic peripheral neuropathy (DPN), which results from the downstream metabolic cascade of longstanding hyperglycemia leading to peripheral nerve injury, is one of the most common complications in diabetic patients. Sensory, autonomic, and motor nerves can all be affected, most frequently in a distal-to-proximal gradient of severity. The most common initial symptoms of DPN are hyperalgesia, dysesthesia and allodynia, and the disease frequently progresses to numbness and hypoalgesia^1^. Though DPN most severely impacts sensory nerves, motor nerve dysfunction can also manifest over the course of the disease^2^. Though the exact mechanism for DPN is unknown, it clearly involves metabolic dysfunction and energy failure in multiple cells within the peripheral nerve and dorsal root ganglion (DRG)^3, 4^.

Growing evidence suggests that lactate is an alternate and effective energy source for the peripheral nerves, and the lactate shuttle between glial and neurons has been demonstrated in the peripheral nervous system (PNS)^5, 6^. The primary transporter for lactate in the PNS, monocarboxylate transporter-1 (MCT1), has been shown to be critical for the response of peripheral nerves to injury^7^, Schwann cell metabolism, and maintenance of sensory nerve myelination during aging^8^. However, the role of MCT1 in the pathogenesis of neurological manifestations of diabetes remains to be explored. Here, we investigate the contribution of MCT1 to the pathogenesis of DPN using the streptozotocin (STZ)-induced diabetes model. We found that deficiency of MCT1 in mice worsens experimental DPN.

## Research Design and Methods

### Animals

All animal experiments were carried out in compliance with the protocols approved by the Johns Hopkins University Institutional Animal Care and Use Committee (IACUC). Wild-type mice aged 8–10 weeks were purchased (C57BL6/J male mice, Jackson Laboratory) and used for the evaluation of MCT1 expression in sciatic nerves and DRG after diabetes induction. Breeding colonies of heterozygous MCT1 null (Het MCT1-null) mice on a C57Bl6 background, obtained from Luc Pellerin and Pierre Magistretti, previously described^7, 9, 10^, were maintained at the Johns Hopkins University. Het MCT1-null mice were used since full knockout of MCT1 is embryonically lethal. Both male and female Het MCT1-null mice and control littermates (wild-type) aged 8–10 weeks were used. Het MCT1-null mice and control littermates develop normally and do not show any notable neurological discrepancies at this age^7^.

### Diabetes induction

Age-matched Het MCT1-null mice and control littermates were used for diabetes induction. As described previously^11, 12^, type-1 diabetes was induced by an intraperitoneal injection of STZ (Sigma-Aldrich; 180 mg/kg body weight) in 0.1 M citrate buffer (pH 4.5). Blood samples were collected from the tail vein three days after the injection, and glycemia was determined with a OneTouch Ultra 2 glucometer (LifeScan Inc., Milpitas, CA). All STZ-injected mice developed hyperglycemia and about 85% of them survived up to the end of the study without any remarkable contribution of any specific genotype.

### RNA preparation and quantitative real-time reverse transcription-PCR

Deeply anesthetized mice were transcardially perfused with 0.1 M PBS to remove the blood, and the sciatic nerves were rapidly dissected. RNA was isolated by an RNeasy Mini Kit (Qiagen), reverse transcribed to cDNA with a High Capacity cDNA Reverse Transcription Kit (Applied Biosystems) and quantified by real-time RT PCR using Taqman probes (Applied Biosystems) for MCT1 (Thermo Fisher Scientific; Catalog # 4351372) or GAPDH (Thermo Fisher Scientific; Catalog # 4352339E) on a StepOne Plus RT-PCR System (Applied Biosystems).

### Nerve conduction studies

Electrophysiologic recordings were performed to measure sensory nerve action potentials (SNAPs) from the tail nerve while maintaining tail temperature at 32–34°C [measured with the Digi-Sense Infrared Thermometer (Model 20250-05)] and compound muscle action potentials (CMAPs) by using a Neurosoft-Evidence 3102evo electromyograph system (Schreiber & Tholen Medizintechnik, Stade, Germany) as described previously^8^.The investigator performing electrophysiologic recordings was blinded to mice genotypes throughout the study.

### Behavioral studies

An experimenter blinded to animal genotype carried out mechanical sensitivity assessment with the von Frey test by frequency method using two calibrated monofilaments (low force, 0.07 g; high force, 0.45 g) and Hargreaves thermal sensitivity test, as described previously^8^. The investigator performing behavioral studies was blinded to mice genotypes throughout the study.

### Intra-epidermal nerve fiber analysis

Analysis of footpads for intra-epidermal nerve fiber density (IENFD) was performed as previously described^8, 13^. The experimenter completing the tissue processing, immunostaining, and analysis was blinded to animal genotype.

### Nerve histology and morphometry

Toluidine blue-stained sections were used for quantification of myelinated axon number, myelinated axon diameter, myelin thickness, or *g* ratio, as described previously^8^. The experimenter performing the morphometric analyses was blinded to animal genotypes.

### Quantification and statistical analysis

Although we did not perform statistical tests to predetermine sample size, our samples sizes are similar to previously published studies in the field. Statistical analyses were performed with GraphPad Prism 8 (GraphPad Software) by using unpaired *t* test with two tails with unequal variance, one-way ANOVA, or two-way ANOVA with post hoc test when required conditions were met. The number of animals per group or independent repeats (*n*), the statistical test used for comparison, and the statistical significance (*p* value) was stated for each figure panel in the respective legend. All data were presented as the mean ± *SEM* unless otherwise noted. Differences in the *p* values of <0.05 were considered statistically significant.

## Results

### MCT1 is reduced early in the sciatic nerve and DRG following induction of STZ diabetes in wild-type mice

To better understand the function of MCT1 in DPN, we first evaluated whether MCT1 expression in the sciatic nerves and DRG is altered in the STZ model of diabetic neuropathy by using quantitative real-time RT-PCR. MCT1 expression was substantially reduced by 2 weeks in both sciatic nerves (**Fig. 1A**) and DRG (**Fig. 1B**), suggesting alterations of lactate pathways and metabolic dysfunctions in diabetic sciatic nerves and DRG.

**Figure 1.**
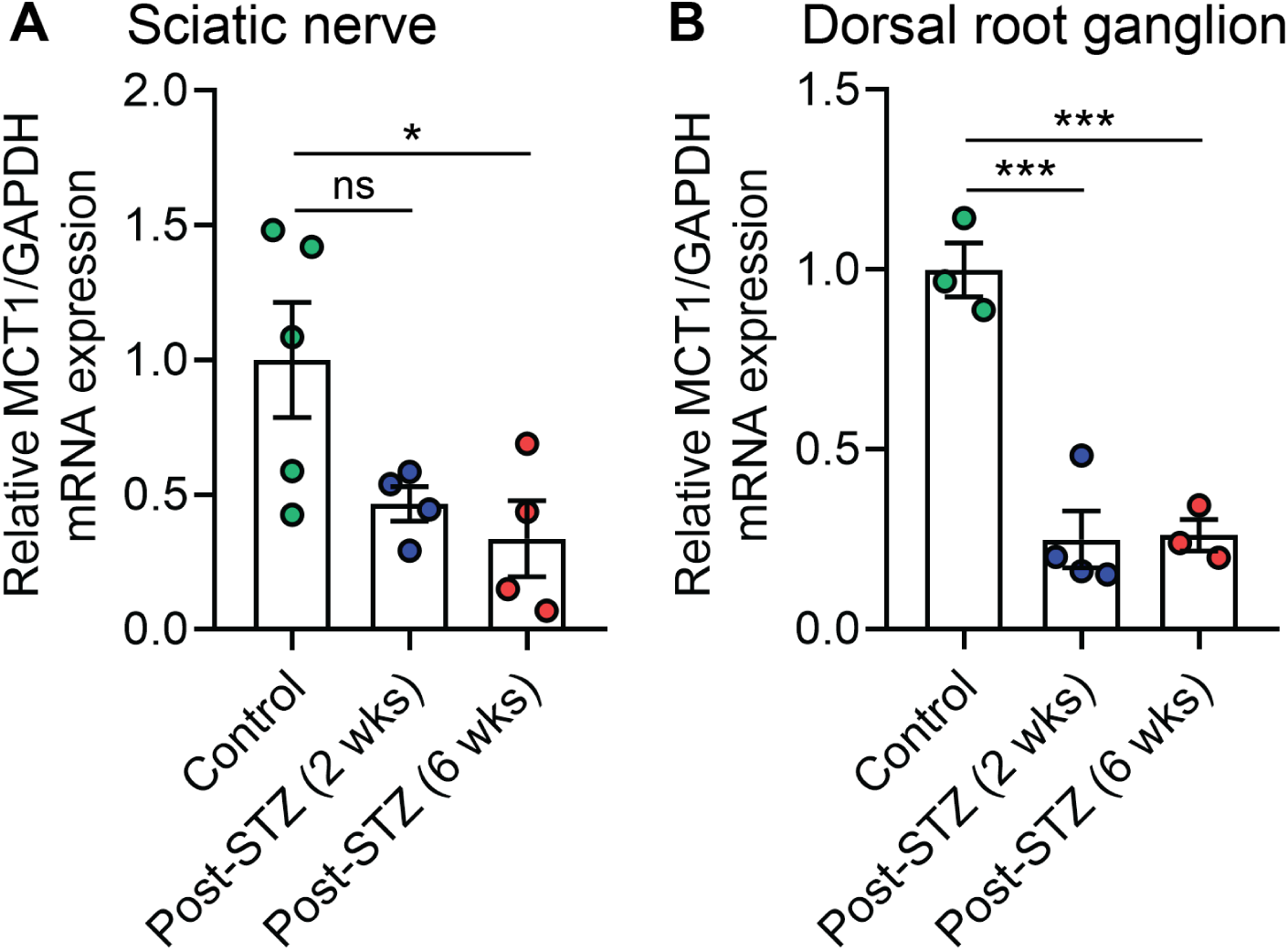
Expression of MCT1 in sciatic nerve and DRG of diabetic mice. The relative expression of MCT1 mRNA in sciatic nerve (**A**) and DRG (**B**) after 2 and 6 weeks of STZ treatment. Levels of mRNA expression are depicted as fold change compared with wild-type mice normalized to their corresponding GAPDH mRNA levels. Mean ± *SEM, n* = 3–5 per group, \**p* < 0.05, \*\*\**p* < 0.001; ns = not significant, one-way ANOVA with Bonferroni’s multiple comparisons test.

### MCT1 reduction does not impact the development of hyperglycemia

The reduction of MCT1 expression in PNS led us to evaluate the impact of reducing MCT1 in diabetes development and pathogenesis of DPN by employing Het MCT1-null mice, which express peripheral nerve MCT1 at approximately 50% of wild-type mice^7^. Despite the reduced expression of MCT1 in peripheral nerve of Het MCT1-null mice, these mice do not show any notable difference in their sensory and motor nerve conductions or mechanical and thermal sensitivities even at the age of 16 weeks compared with their control littermates (**Supplementary Fig. 1**). Since a prior publication showed that these mice do not develop diet-induced obesity and insulin resistance after treatment with high fat diet^10^, we first confirmed that diabetes induction following STZ administration, as measured by hyperglycemia and body weight, was not affected by the reduced expression of MCT1 (**Fig. 2**).

**Figure 2.**
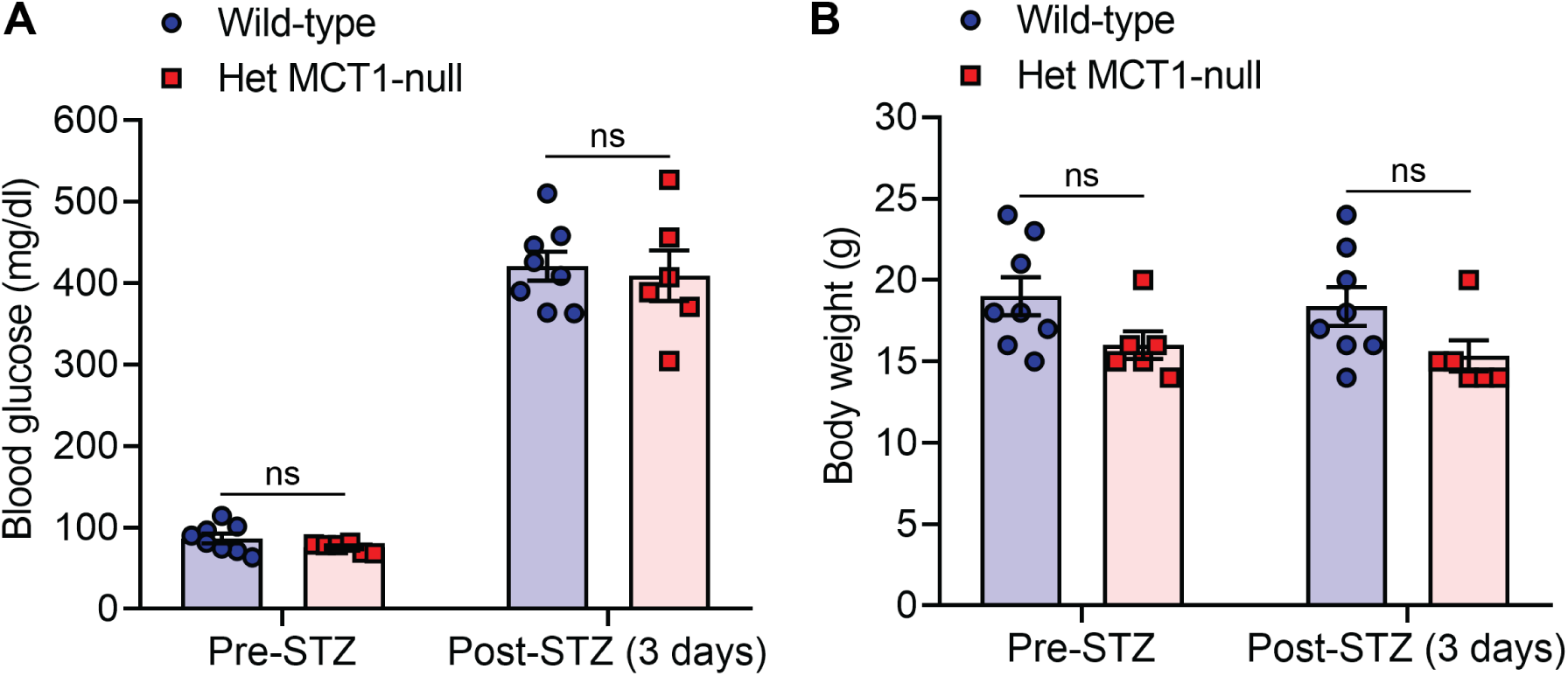
Blood glucose levels and body weight pre- and post-STZ treatment. Blood glucose levels (**A**) and body weight (**B**) were measure before and 3 days after STZ administration. Both groups of mice showed identical extent of hyperglycemia and body weight pre- and post-STZ treatment. Mean ± *SEM, n* = 6–8 per group, ns = not significant, two-way ANOVA with Bonferroni’s multiple comparisons test.

### Reducing MCT1 impairs sensory and motor nerve conductions and reduces mechanical sensitivity

To evaluate the impact of reducing MCT1 on DPN, we measured sensory and motor nerve conduction velocities (NCV) and SNAP and CMAP amplitudes to assess the severity of hyperglycemia-driven nerve damage after diabetes induction. Assessment of NCV is an important indicator of the myelination state of nerves, and SNAP and CMAP amplitudes are indicators of axonal integrity. The NCVs were measured repeatedly from pre-treatment to 9 weeks post-STZ administration (**Fig. 3**, upper panel). Before diabetes induction, mice of both genotypes showed identical sensory and motor NCVs (**Fig. 3A** and **C**) and SNAP and CMAP amplitudes (**Fig. 3B** and **D**), suggesting that reducing MCT1 has no impact on nerve biology, as published previously^7^. However, within 3 weeks of diabetes induction, sensory NCV (**Fig. 3A**) and SNAP (**Fig. 3B**) were significantly decreased in mice having decreased levels of MCT1, compared with control littermates. Furthermore, mice having reduced expression of MCT1 also had impaired motor NCV and CMAP by 6 and 9 weeks, respectively, after diabetes induction. There was no impact of gender on any of these electrophysiological properties. These findings suggest that MCT1 plays an important role in maintenance of sensory and motor nerve function during chronic hyperglycemia.

**Figure 3.**
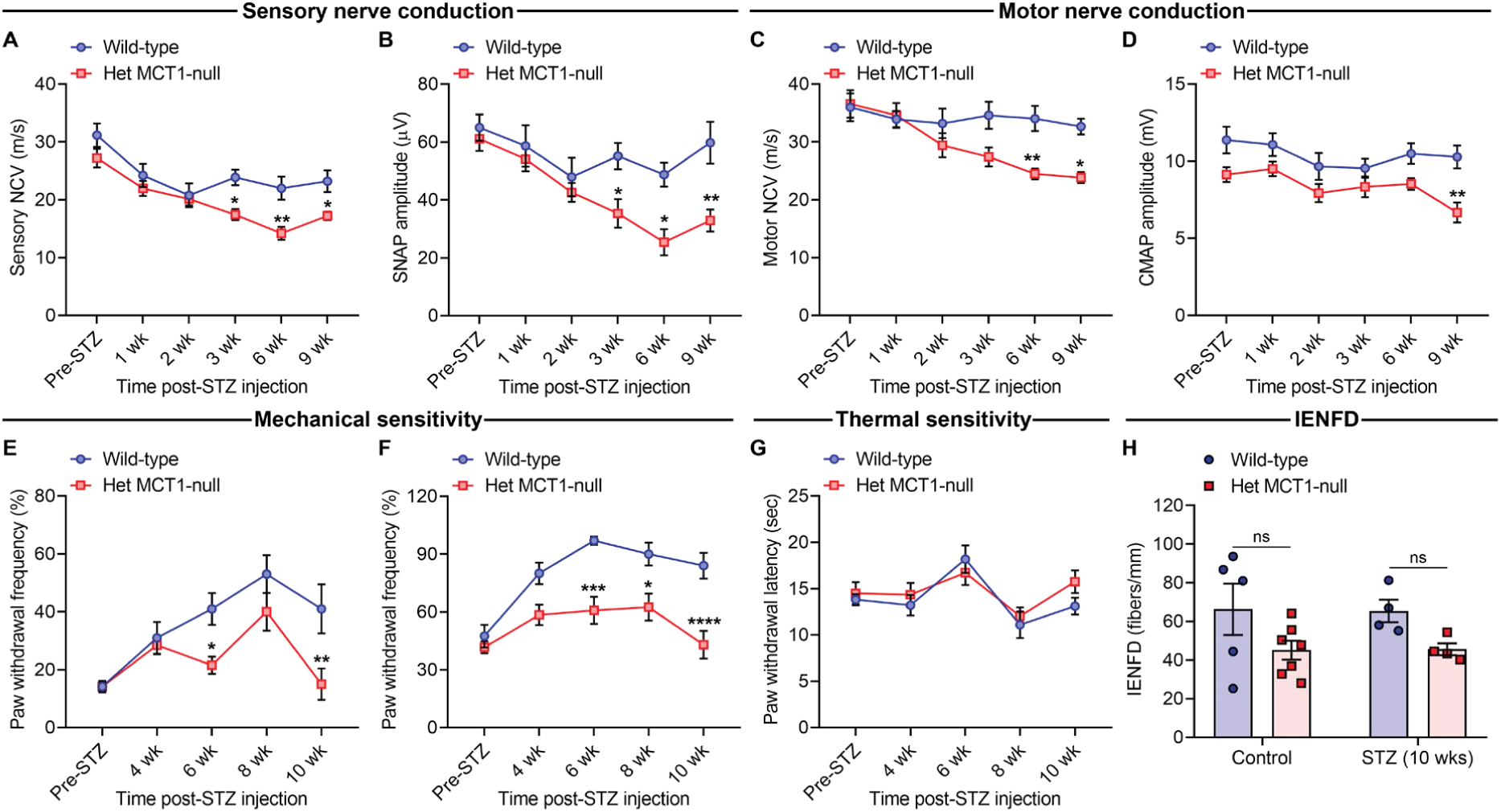
Impact of MCT1 deficiency on nerve conductions and nociceptive behaviors after diabetes induction. Sensory (**A**) and motor (**C**) nerve conduction velocities and SNAP (**B**) and CMAP (**D**) amplitudes in Het MCT1-null mice and control littermates before and after STZ administration. Mean ± *SEM, n* = 7–15 per group, \**p* < 0.05, \*\**p* < 0.01; two-way ANOVA with Bonferroni’s multiple comparisons test. CMAP, compound muscle action potential; NCV, nerve conduction velocity; SNAP, sensory nerve action potential. Paw withdrawal frequency to mechanical stimulation by calibrated von Frey monofilaments of forces 0.07 g (**E**) and 0.45 g (**F**) and paw withdrawal latency (**G**) to thermal stimulation by radiant paw-heating assay in Het MCT1-null mice and control littermates before and after STZ administration. Current set at baseline level: 20%, 10–12 s; cut off time; 30 s. Mean ± *SEM, n* = 6–13 per group, \**p* < 0.05; \*\**p* <0.01; \*\*\**p* <0.001; \*\*\*\**p* <0.0001, two-way ANOVA with Bonferroni’s multiple comparisons test. (**H**) IENFD obtained from the footpads of control or diabetic mice at 10 weeks after STZ administration following immunohistochemical staining for PGP9.5. Mean ± *SEM, n* = 4–7 per group, ns = not significant, two-way ANOVA with Bonferroni’s multiple comparisons test. IENFD, intraepidermal nerve fiber density

Impaired nerve electrophysiology in diabetic mice with reduced MCT1 expression led us to investigate if there were any nociceptive behavioral deficits in these mice. The behavioral responses to repetitive punctate mechanical stimulation and noxious thermal stimulation were assessed with the von Frey test by the frequency method and the Hargreaves test, respectively (**Fig. 3E-G**). Mice with or without reduced MCT1 showed no difference in mechanical and thermal sensitives before STZ administration. However, mice with reduced MCT1 expression had decreased mechanical sensitivity within 6 weeks of diabetes induction, as indicated by decreased paw withdrawal frequency to repetitive high force (0.45 g) von Frey filament stimulation (**Fig. 3F**). This reduced mechanical sensitivity in Het MCT1-null mice is in contrast to wild-type mice that show increased sensitivity, or allodynia, over this same time period (**Fig. 3E** and **F**). In contrast to mechanical sensitivity, no alteration in paw withdrawal latency to noxious thermal stimulation was observed in Het MCT1-null mice after diabetes induction (**Fig. 3G**). Furthermore, the unchanged thermal sensitivity after diabetes induction was further supported by unchanged IENFD, as measured by PGP9.5-immunoreactive nerve counts normalized to epidermal area, which measures unmyelinated nociceptive nerve fibers in the skin (**Fig. 3H**). We did not see any remarkable impact of gender on these behaviors or histology. These findings suggest that MCT1 reduction impairs mechanical, but not thermal, sensitivity in diabetes mice.

### Reduction in MCT1 results in sural nerve demyelination after diabetes induction

The nerve conduction velocity (NCV) substantially depends on the myelination status of nerve. Thus, we investigated the morphology of sural nerves isolated from Het MCT1-null and control littermate diabetic mice. Consistent with the electrophysiologic recordings of sensory NCV, we observed significant demyelination (**Fig. 4A**) due to MCT1 reduction, measured both by *g* ratio (**Fig. 4B** and **C**) and myelin thickness (**Fig. 4D**). However, unlike change in SNAP amplitude of tail sensory nerve, there was no change in the number of myelinated axons per sural nerve (**Fig. 4E**), suggesting that MCT1 plays a critical role in myelin maintenance, but not axonal maintenance, during DPN pathogenesis.

**Figure 4.**
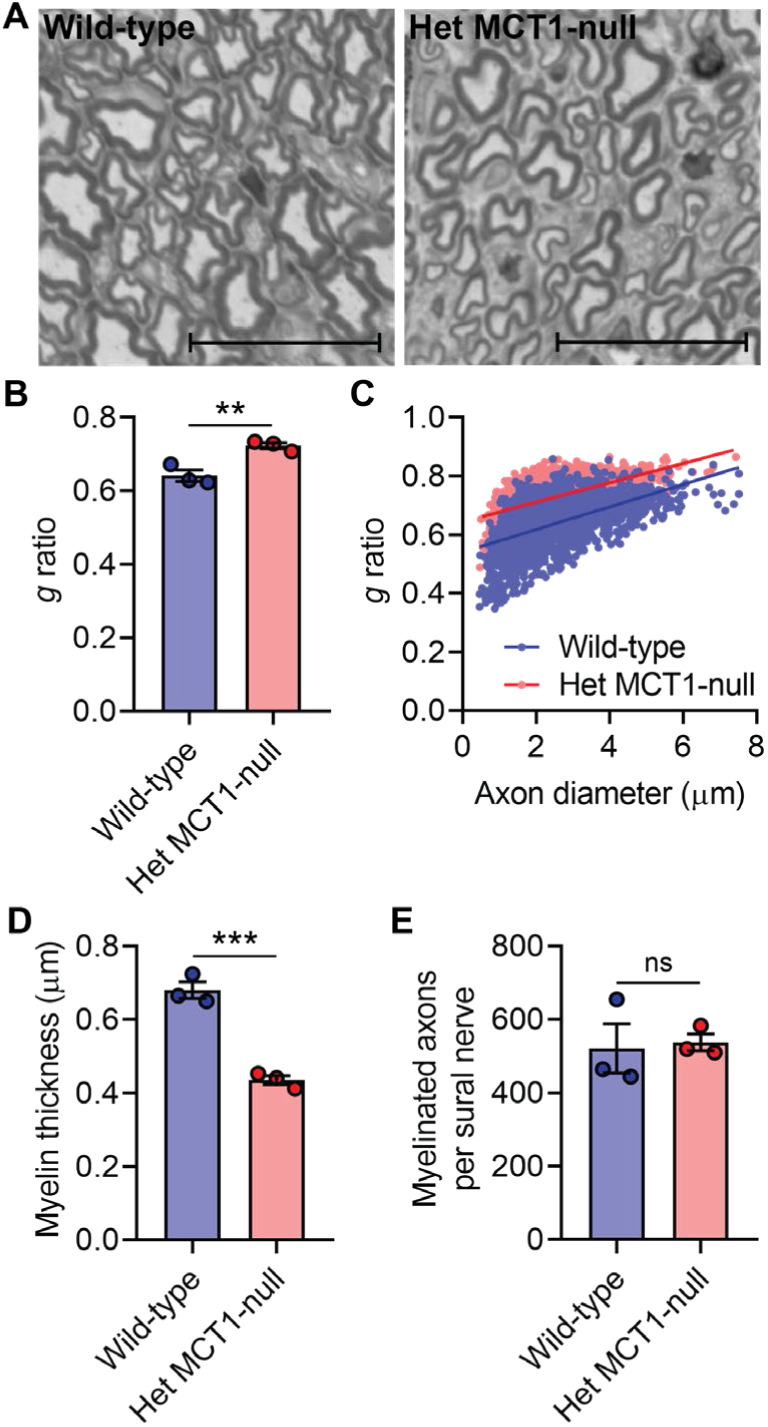
Impact of reduced MCT1 on sural nerve myelination and integrity in diabetes. (**A**) Light microscope photomicrographs of toluidine blue-stained sections of sural nerves from Het MCT1-null mice and control littermates after 10 weeks of STZ treatment. These images were analyzed for *g* ratio (**B**), scatter plot graph displaying *g* ratio (*y*-axis) in relation to axon diameter (*x*-axis) of individual fiber (**C**), myelin thickness (**D**), and myelinated axon counts (**E**). Mean ± *SEM, n* = 3 per group, \*\**p* < 0.01; \*\*\**p* < .001; ns = not significant, unpaired *t* test. Scale bar, 20 μm. The *g* ratio between wild-type (blue line) and Het MCT1-null (red line) mice (**C**) was significantly different (*p* < 0.0001; *t* = 30.75, *df* = 3,177, unpaired *t* test).

## Discussion

In this study, we evaluate, for the first time, the potential role of MCT1 in the pathogenesis of DPN. This study suggests that chronic hyperglycemia decreases the expression of MCT1 in PNS, and mice with reduced expression of MCT1 exhibit progressive NCV slowing and reduced SNAP as well as CMAP amplitudes, electrophysiological features of DPN in animal models and humans^14, 15^. In addition, this study demonstrates that these electrophysiological deficits are due to demyelination. This study further demonstrates that mice with reduced expression of MCT1 develop depressed sensitivity to mechanical stimulation. Though likely reflecting increased numbness, this reduced sensitivity to mechanical stimulation is not necessarily detrimental, since it appears to be lowering the development of mechanical allodynia.

Converging evidence suggests that DPN is at least partly due to energy failure in the peripheral nerve^3, 4^. MCT1 is the predominant lactate transporter throughout the body^16^, and its expression is altered in adipocytes, muscle, and brain^17-19^ of patients and animal models with diabetes. Despite its critical role in nerve regeneration after injury^7^ and nerve myelination during aging ^8^, MCT1 has not previously been investigated in DPN. This study shows that in STZ-treated wild-type mice, MCT1 is reduced early in both sciatic nerves and DRG, prior to any axon or Schwann cell degeneration^12, 20^, suggesting a possible contribution to metabolic dysfunction and resultant energy failure in diabetic sciatic nerves and DRG.

MCT1 deficiency accelerates DPN and sensory loss in Het MCT1-null mice, and the worsening of neuropathy occurs without any change in the degree of hyperglycemia, suggesting that MCT1 does not contribute to these abnormalities directly by modulating the extent of hyperglycemia. MCT1 potentially modulates the metabolism of diverse cells associated with the pathophysiology of DPN. The increased demyelination in sural nerves from diabetic Het MCT1-null mice suggests that MCT1 is critical for the maintenance of myelin integrity during DPN, potentially due to an important role in Schwann cells. This finding is consistent with our recent study, which demonstrated a role for Schwann cell-specific MCT1 in the maintenance of sensory nerve myelination during aging by modulating lipid metabolism in peripheral nerves^8^. Importantly, however, the changes in myelination following STZ treatment of Het MCT1-null mice is not merely due to aging, since sensory NCV reductions were seen in STZ-treated mice before 3 months of age, which is prior to the previously published hypomyelination observed with aging. Additionally, there were no changes in SNAP amplitude, motor NCV, or motor CMAP amplitude, at any timepoint, following Schwann cell-selective knockdown of MCT1^8^.

DPN in patients preferentially targets distal components of sensory axons, and only later, and usually to a lesser extent, motor axons^2, 21^. In this study, reduced expression of MCT1 causes NCV deficits in sensory nerves within 3 weeks and motor nerves within 6 weeks of diabetes induction. Furthermore, Het MCT1-null mice have progressive declines in SNAP and CMAP amplitudes without any remarkable change in the number of myelinated sural nerve axons. There are two possible explanations for these apparently contradictory findings. First, severe demyelination may be causing conduction block, in which action potentials being conducted along a demyelinated nerve are unable to propagate due to a large gap in myelin^22^. Second, since the axon degeneration in DPN is thought to be primarily a “dying back” process that impacts the most distal components of the nerve^23^, perhaps the sural nerve counts are unchanged because they are measured from more proximal nerve segments than those measured by tail SNAPs. Further confirmation of distal sensory nerve injury comes from behavioral testing, as Het MCT1-null mice treated with STZ have reduced response to mechanosensitivity with von Frey monofilaments.

In summary, our findings demonstrate that MCT1 plays an important role in DPN pathogenesis. MCT1 expression is reduced in both sciatic nerve and DRG following STZ-induced diabetes, and reducing MCT1 worsens DPN in this model of type 1 diabetes. The mechanism by which MCT1 contributes to DPN is not clear, but may be due to reduced capacity of cells without MCT1 to process elevated glucose levels, which would be expected to generate higher amounts of lactate, or impaired PNS cell metabolism leading to deficits in metabolic support to Schwann cells or neurons. Regardless of mechanism, our results suggest that manipulating this transporter may be a potential new avenue for DPN treatments. Additionally, the results of our paper suggest that clinical development of MCT1 inhibitors, either as immunosuppressive agents^24^ or chemotherapies^25^, should proceed cautiously, as there may be unexpected side effects in patients with diabetes.

## Acknowledgements

The authors would like to thank Ms. Kimberly Brown and the Johns Hopkins Neurology Electron Microscopy Core for their assistance in processing embedded nerve tissue for toluidine blue staining. We would also like to thank the Pain Research Core funded by the Blaustein Fund and the Neurosurgery Pain Research Institute at The Johns Hopkins University for providing facilities for behavioral studies. Financial support was provided by NIH-NS086818-01 (B.M.M.). B.M.M. is the guarantor of this work and, as such, had full access to all the date in the study and takes responsibility for the integrity of the data and the accuracy of the data analysis.

## Author Contributions

M.K.J. conceived and performed experiments, analyzed data, and wrote the manuscript. X.H.A., F.Y., and Y.L. performed some of the experiments and/or data analysis. M.J.P. and L.P. assisted with resources. B.M.M. secured funding, conceived and supervised the study, analyzed data, and wrote the manuscript.

## Conflict of Interest

The authors declare no competing financial interests.

## Figures

**Supplementary Figure 1.**
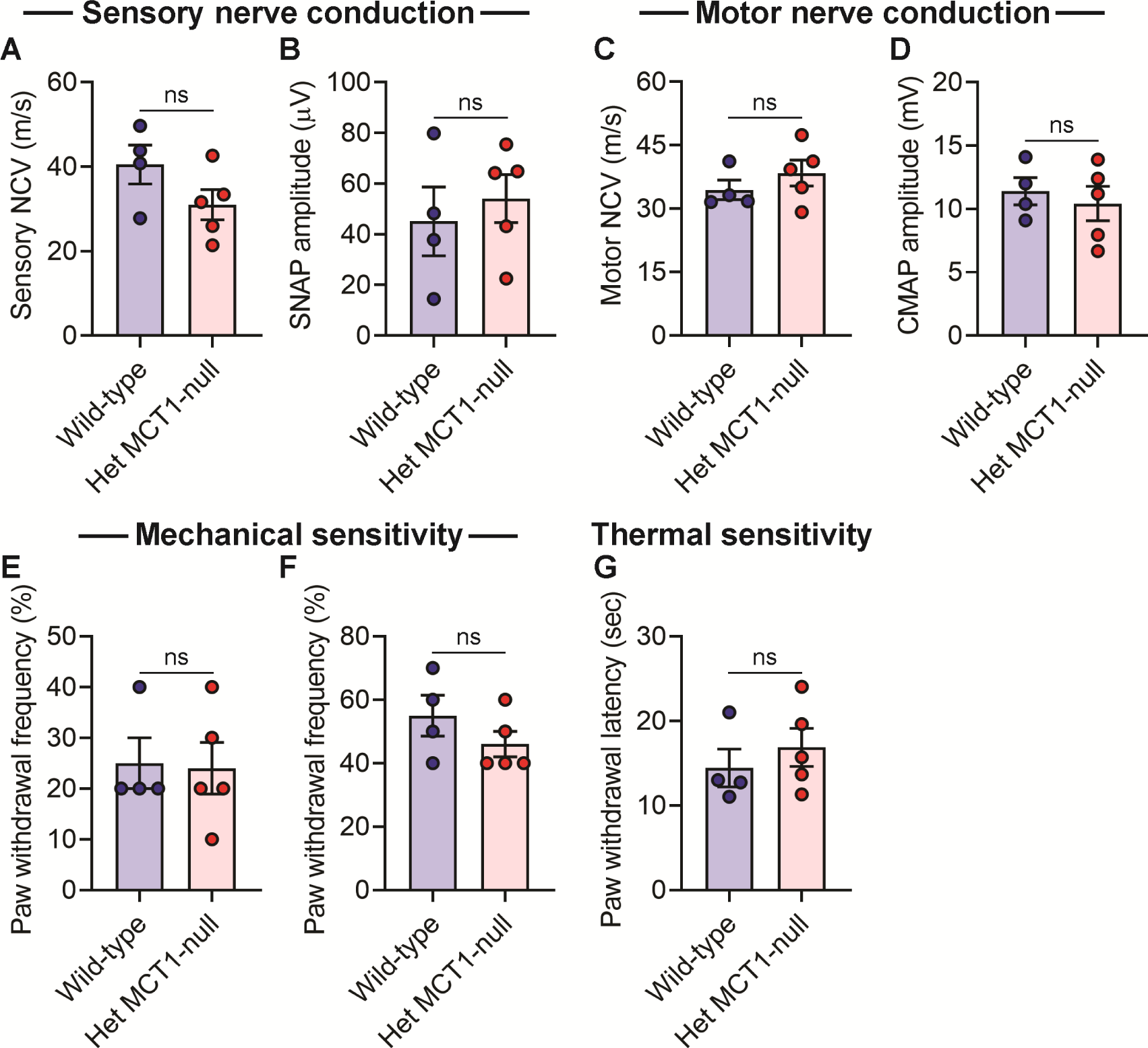
Impact of MCT1 deficiency on nerve conductions and nociceptive behaviors in 16-week-old non-diabetic mice. Sensory (**A**) and motor (**C**) nerve conduction velocities and SNAP (**B**) and CMAP (**D**) amplitudes in 16-week-old Het MCT1-null mice and control littermates without any treatment. Paw withdrawal frequency to mechanical stimulation by calibrated von Frey monofilaments of forces 0.07 g (**E**) and 0.45 g (**F**) and paw withdrawal latency (**G**) to thermal stimulation by radiant paw-heating assay in Het MCT1-null mice and control littermates. Current set at baseline level: 20%, 10–12 s; cut off time; 30 s. Mean ± *SEM, n* = 4–5 per group, ns = not significant, unpaired *t* test. CMAP, compound muscle action potential; NCV, nerve conduction velocity; SNAP, sensory nerve action potential.

